# Inter-domain Horizontal Gene Transfer of Nickel-binding Superoxide Dismutase

**DOI:** 10.1101/2021.01.12.426412

**Authors:** Kevin M. Sutherland, Lewis M. Ward, Chloé-Rose Colombero, David T. Johnston

## Abstract

The ability of aerobic microorganisms to regulate internal and external concentrations of the reactive oxygen species (ROS) superoxide directly influences the health and viability of cells. Superoxide dismutases (SODs) are the primary regulatory enzymes that are used by microorganisms to degrade superoxide. SOD is not one, but three separate, non-homologous enzymes that perform the same function. Thus, the evolutionary history of genes encoding for different SOD enzymes is one of convergent evolution, which reflects environmental selection brought about by an oxygenated atmosphere, changes in metal availability, and opportunistic horizontal gene transfer (HGT). In this study we examine the phylogenetic history of the protein sequence encoding for the nickel-binding metalloform of the SOD enzyme (SodN). A comparison of organismal and SodN protein phylogenetic trees reveals several instances of HGT, including multiple inter-domain transfers of the *sodN* gene from the bacterial domain to the archaeal domain. Nearly half of the archaeal members with *sodN* live in the photic zone of the marine water column. The *sodN* gene is widespread and characterized by apparent vertical gene transfer in some sediment-associated lineages within the Actinobacteriota (Actinobacteria) and Chloroflexota (Chloroflexi) phyla, suggesting the ancestral *sodN* likely originated in one of these clades before expanding its taxonomic and biogeographic distribution to additional microbial groups in the surface ocean in response to decreasing iron availability. In addition to decreasing iron quotas, nickel-binding SOD has the added benefit of withstanding high reactant and product ROS concentrations without damaging the enzyme, making it particularly well suited for the modern surface ocean.

## Introduction

Molecular oxygen in the ground state is a diradical. The presence of these two unpaired electrons in parallel spin makes O_2_ susceptible to univalent reduction, which will lead to the formation of reactive oxygen species (ROS) (Fridovich, 1978). When O_2_ interacts with an electron donor such as a reduced metal or organic carbon and univalent reduction occurs, superoxide (O_2_^•−^) is formed. This reaction occurs in association with the core metabolic and physiological functions of microorganisms. Superoxide is produced at photosystem I during photosynthesis (Asada, 2006), at multiple locations along the electron transport chain in aerobic respiration (Larosa & Remacle, 2018), and on the outer membrane of cells for a range of purposes including nutrient acquisition (Rose et al., 2008), cell signaling (Buetler et al., 2004), redox homeostasis (Diaz et al., 2019; Yuasa et al., 2020), and cell growth and proliferation (Carlioz & Touati, 1986; Oda et al., 1995; Saran, 2003). Superoxide is a highly reactive molecule that typically has a lifetime on the order of a few minutes under physiological conditions (Diaz et al., 2013). It can further act as an oxidant or reductant in a wide range of consequential biogeochemical reactions involving nutrients, metals, organic carbon, and nitrogen species.

The evolution of oxygenic photosynthesis and subsequent Great Oxygenation Event (~2.3-2.4 Gya) greatly increased the energy available to fuel microbial metabolisms (Fischer et al., 2016; Ward et al., 2019). However, increasing oxygen levels is in many ways a double-edged sword. The potential to produce superoxide and other ROS would have increased in parallel with atmospheric oxygen levels, which can have a myriad of detrimental effects on cells (Taverne et al., 2018). The reactivity of superoxide with a wide range of organic moieties and metal cofactors can damage various cell components, including lipid membrane peroxidation and iron release from iron-sulfur clusters (Longo et al., 1999; Radi et al., 1991). Superoxide production can precede the formation of hydroxyl radical, which itself can have devastating effects on biomolecules (Buxton et al., 1988; Oda et al., 1992). The ability to manage and mitigate the deleterious consequences of superoxide and its downstream effects and reaction products is an inescapable challenge of life evolving in the presence of molecular oxygen.

Superoxide dismutase is the primary biological tool that microorganisms use to degrade superoxide, both within the cell membrane and in the environment. The presence of at least one copy of superoxide dismutase in the genome of all extant aerobic organisms (and many anaerobes) underscores its necessity in coping with reactive oxygen (Brioukhanov & Netrusov, 2004). The name “superoxide dismutase” is used as a catch-all term to describe four different metalloforms of three distinct enzyme families that have no known homology. These four metalloforms include manganese (Mn) (gene: *sodA*, protein: SodA, enzyme: MnSOD), iron (Fe) (gene: *sodB*, protein: SodB, enzyme: FeSOD), copper and zinc (CuZn) (gene: *sodC*, protein: SodC, enzyme: CuZnSOD), and nickel (Ni) (gene: *sodN*, protein: SodN, enzyme: NiSOD)(Dupont et al., 2008; Wolfe-Simon et al., 2005). Only the Mn- and Fe-binding metalloforms have demonstrable homology (including a cambialistic isoform that can function with either Fe or Mn (Lancaster et al., 2004)), thus, superoxide dismutase is likely an example of convergent evolution in which life reinvented the wheel three separate times (No, 2017).

The concentration of oxygen, its influence on metal redox state, and the availability of each metal cofactor in the environment all present important selective pressures on determining the distribution of these different SOD-encoding genes spatially and throughout Earth History. While the true antiquity of these three enzyme families is not known with certainty, the relative ages have been inferred from a combination of phylogenetic arguments and likely environmental conditions on the ancient surface Earth. The FeSOD/MnSOD enzyme family is thought to be the most ancient of these three enzyme families (Hatchikian & Henry, 1977; Yost & Fridovich, 1973). This notion is evidenced by the observation that *sodA/sodB* is phylogenetically wide-spread, and because low atmospheric oxygen in the early Earth would have favored high concentrations of dissolved Fe^2+^ and Mn^2+^ (Wolfe-Simon et al., 2005). Conversely, CuZnSOD, thought to be the youngest of the three enzyme families, is widespread in higher plants and animals, present in some bacteria, and absent from archaea and protists (Case, 2017). This metalloform of SOD can also be localized to the cytoplasm, the nucleus, mitochondrial intermembrane space, chloroplasts, and extracellular space (Case, 2017; Weisiger & Fridovich, 1973). The presumed younger age of the CuZnSOD is supported by its relative scarcity among microorganisms, and the fact that Cu and Zn would have both been highly insoluble in an low-oxygen world (Case, 2017; Saito et al., 2003; Wolfe-Simon et al., 2005).

Intermediate in antiquity to the MnSOD/FeSOD and CuZnSOD enzymes, and the primary focus of this study, is Ni-binding superoxide dismutase. NiSOD is the most recently discovered of these three enzyme families (discovered in 1996), and has been the subject of less study than the other two enzyme families (Youn et al., 1996). The evolution and proliferation of NiSOD is thought to coincide with decreasing iron availability resulting from ocean oxygenation (Wolfe-Simon et al., 2005). In fact, NiSOD is the most common metalloform of SOD in modern marine bacteria (Sheng et al., 2014). The abundance of NiSOD among marine Cyanobacteria has been used to suggest that NiSOD may have originated among marine phototrophs (Case, 2017; Wolfe-Simon et al., 2005). However, more recent work suggests that *sodN* originated within the Actinobacteriota clade before appearing in other clades through a combination of vertical and horizontal gene transfer (Schmidt et al., 2009). Currently, *sodN* is thought to be present primarily in bacterial clades, however, some instances of *sodN* in photosynthetic eukaryotes have been identified (Cuvelier et al., 2010). To date, *sodN* has not been identified in archaea (Sheng et al., 2014).

Given the ever-expanding availability of quality genomic data and analysis tools, we revisit the phylogeny of the NiSOD protein to better understand its evolutionary history and its distribution across the tree of life. We examine the role of vertical and horizontal gene transfer in the controlling the phylogeny of NiSOD sequences. With this information, we aim to better understand the environmental pressures that select for NiSOD over other SOD genes and contextualize the history of this gene within the broader scope of global biogeochemical cycles.

## Methods

A reference database was constructed containing a single high-quality genome from every genus of bacteria and archaea for which one is available based on the GTDB database (Parks et al., 2018, 2020). A list of all genomes included in the GTDB database was downloaded from the GTDB servers (https://data.ace.uq.edu.au/public/gtdb/data/releases/). Genomes that did not meet current metrics for high quality (e.g. completeness <90% or contamination >5%, (Bowers et al., 2017)) were removed. This list was finally dereplicated to select a single genome of the highest available completeness from every genus. Genomes were then downloaded from the NCBI WGS and Genbank databases. This genome database was used to search for homologs of SodN and other proteins using the tblastn function of BLAST+ (Camacho et al., 2009). For organismal phylogenies, concatenated ribosomal protein phylogenies following methods from Hug et al. (2016) (Hug et al., 2016). Protein sequences were aligned with MUSCLE (Edgar, 2004) and phylogenetic trees were produced using RAxML v.8.2.12 (Stamatakis, 2014) on the Cipres Science Gateway (Miller et al., 2010). Node support was calculated using transfer bootstrap support with BOOSTER (Lemoine et al., 2018). Trees were visualized using the Interactive Tree of Life Viewer (Letunic & Bork, 2016). Histories of horizontal gene transfer were assessed via incongruence of organismal and metabolic protein phylogenies as described previously (e.g. (Doolittle, 1986; Ward et al., 2020; Ward & Shih, 2020)). Assessment of genomic neighborhood of sodN genes was performed on the RAST platform (Aziz et al., 2008). In order to verify that NiSOD presence was authentic and not a case of false positives (i.e. gene presence in metagenome-assembled genomes due to contamination, e.g. (Ward et al., 2018)), we repeated analyses with a more stringent quality cutoff of a maximum of 3.5% contamination as determined by CheckM (Parks et al., 2015).

## Results

We have produced the most complete phylogeny of NiSOD proteins available. The phylogenies of the organisms identified in this study and their respective SodN protein sequences are presented in Figures 1 and 2, respectively. This SodN phylogeny expands substantially on the small collection of characterized NiSOD-producing isolates to include many lineages known only or primarily from metagenome-assembled genomes (e.g. Patescibacteria) as well as cultured organisms in which the presence of NiSOD was previously undescribed (e.g. diverse Proteobacteria and Actinobacteriota). This more comprehensive view of the distribution of NiSOD provides a much more complete context for interpreting the evolutionary history of this enzyme family. At present, no homologous outgroup to the NiSOD protein family is known, and as a result it impossible to robustly root the NiSOD phylogeny. For ease of viewing Figure 2 is shown with an arbitrary midpoint root, but this is not intended to imply directionality of evolution. Nonetheless, our results provide valuable understanding to evolutionary relatedness between NiSOD sequences even if we are unable to pinpoint the origin of this enzyme family.

**Fig 1:**
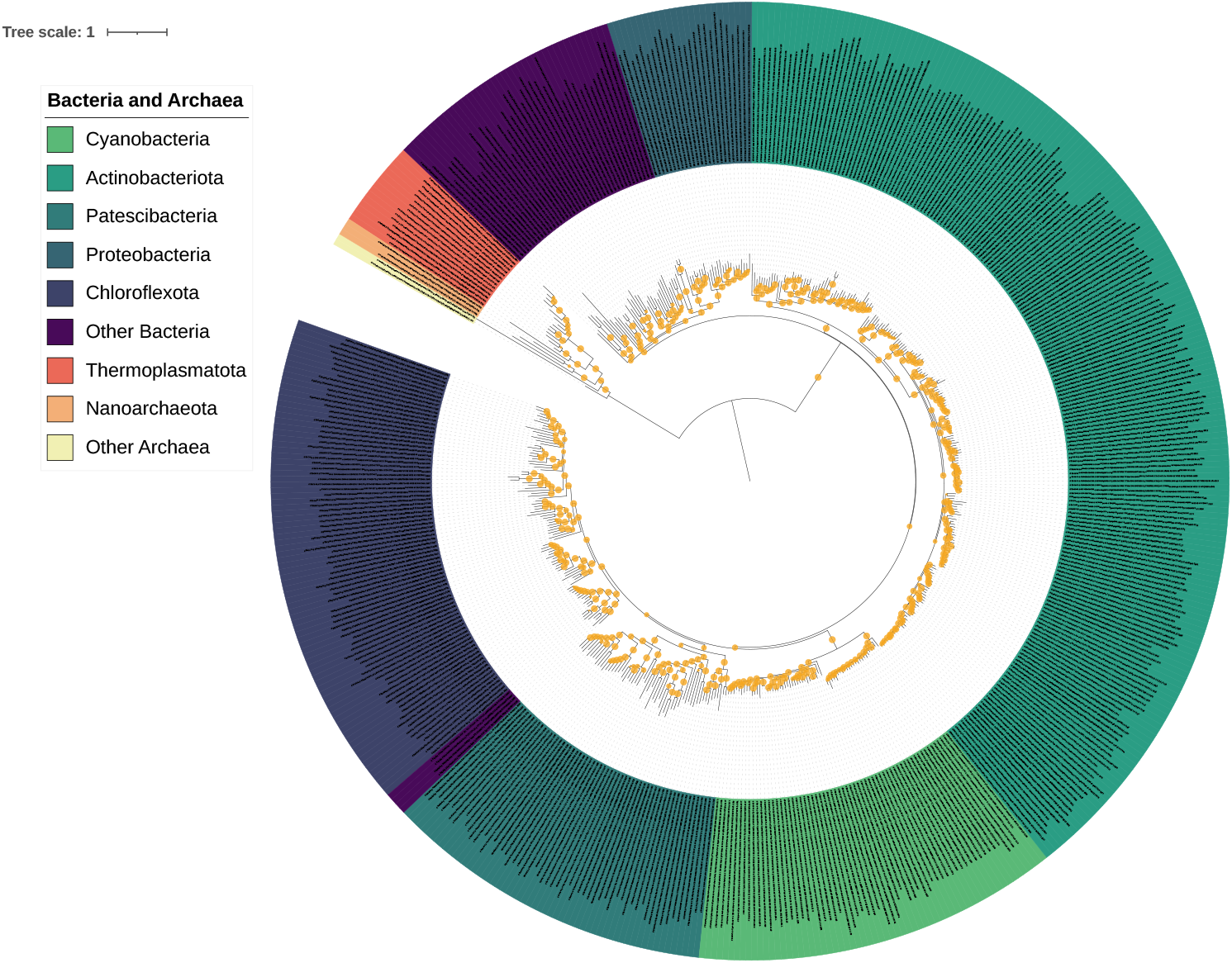
Organismal phylogenetic tree determined using concatenated ribosomal protein sequences of organisms from the Genome Taxonomy Database that contain a *sodN* gene. We note that there are slightly fewer organisms in the organismal tree than in the SodN protein tree (Figure 2), this is due to either low completeness or poorly aligned concatenated ribosomal sequences. See methods for a complete description of tree production and quality cutoff metrics.

**Fig 2:**
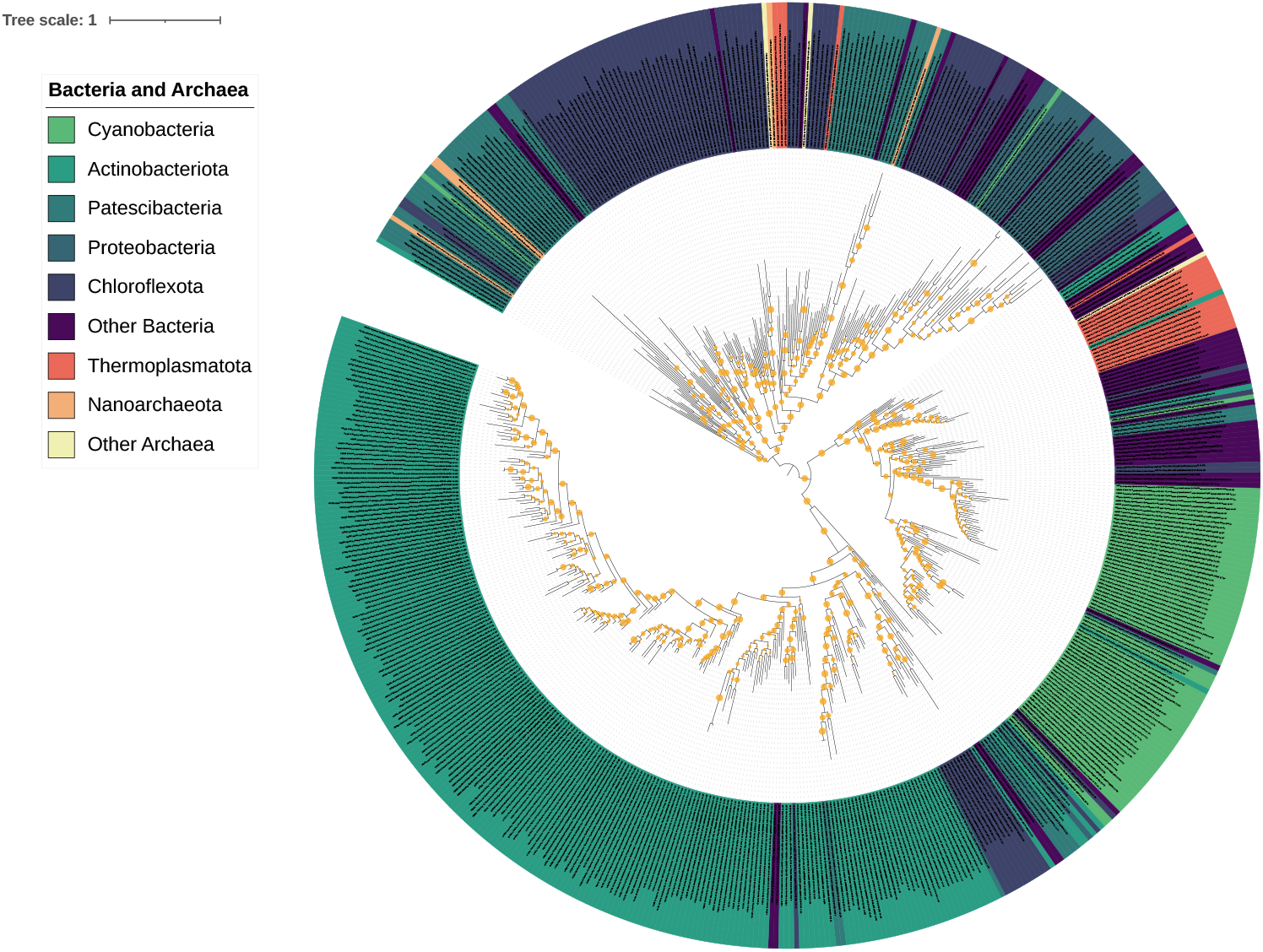
SodN protein sequence tree for organisms containing at least one copy of the *sodN* gene in the Genome Taxonomy Database.

The presence of a small number of deep clades is readily visible in the NiSOD phylogeny. In particular, much of the diversity of NiSOD sequences belongs to a large clade made up primarily of sequences from Actinobacteriota, but also containing large clusters of sequences from Cyanobacteria and smaller lineages from other organisms (bottom half of Figure 2). The remainder of NiSOD diversity belongs to a somewhat smaller but more diverse clade made up largely of Chloroflexota sequences interspersed with sequences from Patescibacteria, Proteobacteria, and other organisms (top half of Figure 2). While each of these clades contains sequences from multiple bacterial phyla (and, surprisingly, multiple archaeal phyla), these clades are made up of a plurality of and bounded by lineages of Actinobacterota and Chloroflexota, respectively. With few exceptions (e.g. Cyanobacteria), NiSOD sequences from a particular phylum are polyphyletic, scattered in multiple lineages separated by many sequences from other phyla. Moreover, the topology of phylogenetic relationships both within and between phyla are largely incongruent with organismal relationships, indicating many instances of interphylum horizontal gene transfer.

In contrast to previous claims (Sheng et al., 2014), we found that NiSOD are not absent from the archaeal domain. In fact, several clades of archaeal NiSOD exist, scattered throughout the enzyme phylogeny (Figure 2). The lack of reciprocal monophyly between bacterial and archaeal NiSOD sequences, and the incongruence between organismal and protein phylogenies (Figure 1, Figure 2), strongly suggests that the distribution of NiSOD genes throughout the tree of life is the result of multiple instances of interdomain horizontal gene transfer.

## Discussion

Horizontal gene transfer (HGT) is commonplace in the microbial world. It is invoked to explain the distribution of a wide variety of genes that play a role in all aspects of metabolism and physiology. Evidence of inter-domain horizontal gene transfer, however, is much less commonplace. Generally speaking, the frequency of horizontal gene transfer between prokaryotes is thought to increase with increasing genetic similarity, meaning inter-domain horizontal gene transfer should be the rarest genetic transfer (Andam & Gogarten, 2011). In fact, clear instances of horizontal gene transfer between archaea and bacteria are fleeting and thought to be quite rare (Avni & Snir, 2020). It is clear from this study that NiSOD has indeed undergone multiple instances of horizontal gene transfer, including several instances of transfer from the bacterial domain to the archaeal domain.

NiSOD sequences are distributed throughout the bacterial domain, with most protein diversity distributed between two large clades made up primarily of Actinobacteriota and Chloroflexota. Substantial diversity of sequences derived from other bacterial and archaeal taxa appears interspersed within these two clades, apparently reflecting a long history of horizontal gene transfer within and between deep taxonomic groups including at the phylum and domain level. While support values for some shallow nodes are relatively poor (e.g. within closely related clades of Actinobacteriota sequences), support values for deep nodes including those between archaeal and bacterial lineages are robust, providing strong confidence in interpretations of horizontal gene transfer. Because of our stringent quality cutoff for metagenome-assembled genomes (MAGs), we can confidently assert that the organismal source of NiSOD proteins is accurate, and does not reflect contamination or incorrect recruitment of sequences into MAGs, further strengthening interpretations of HGT.

Many apparent instances of HGT of NiSOD—including the bulk of interdomain transfer events—appear to have occurred between taxa that tend to inhabit similar niches within the same environment. For example, most Chloroflexota NiSOD sequences are from members of the Dehalocoiccoidia class. Members of Dehalocoiccoidia are abundant and diverse in marine sediment, terrestrial soil, and groundwater primarily as anaerobic heterotrophs (Taş et al., 2010; Wasmund et al., 2016). Nested within the Chloroflexota NiSOD sequences are at least three clusters of sequences from the Patescibacteria. Nested even deeper within Patescibacteria NiSOD clades are a few sequences derived from archaea including the Woesearchaea. Both Patescibacteria and Woesearchaea are thought to exploit similar strategies in the terrestrial subsurface as symbionts with highly reduced genomes and cell sizes (Brown et al., 2015; Castelle et al., 2015). This suggests the possibility that communities of predominantly anaerobic heterotrophs in terrestrial sediment may have exchanged NiSOD genes repeatedly over geologic time, likely as an adaptation to tolerate trace amounts of oxidants. This appears to have resulted in HGT of NiSOD from Chloroflexota to Patescibacteria in at least three events, followed by two transfers from Patescibacteria to Woesearchaea. In this case, it appears that colocalization and overlapping metabolic strategies has enabled repeated HGT between lineages that have diverged on the superphylum and even the domain level.

Similarly, interdomain HGT of NiSOD appears to have occurred between organisms in shared environments with similar life strategies. An example of this is seen within the Actinobacteriota clade of NiSOD sequences. Here, NiSOD sequences derived from the archaeal Poseidoniales clade appear nested within a deeper group of bacterial Marinisomata sequences. Both Marinisomata and Poseidoniales are primarily known to exist as water column heterotrophs, with Poseidoniales primarily restricted to the photic zone while Marinisomata exist throughout the water column (Huang & Wang, 2020; Rinke et al., 2019). This suggests a possible route of transmission of NiSOD genes from marine sediment Actinobacteriota, to water column Marinisomata, and ultimately to surface ocean Poseidoniales. A similar transmission process through different taxa moving up the water column may have led to the acquisition of NiSOD by Cyanobacteria. However, Cyanobacteria and Poseidoniales NiSOD sequences are not closely related, suggesting that NiSOD sequences were introduced into the photic zone multiple times from multiple lineages. This repeated, directional migration of NiSOD genes through the environment, rather than extensive sharing of a single set of NiSOD genes simply within the photic zone, highlights that the successful horizontal transfer of genes can require more than simple colocalization of organisms and that multiple routes of transfer can be occurring into and within an environment at any given time.

There are two likely candidates for the clade in which *sodN* originated: Actinobacteriota and Chloroflexota. The *sodN* gene is widely distributed across both clades, and the phylogeny of the SodN protein largely resembles the organismal tree of these two clades. In the absence of a known outgroup, it is not formally possible to root SodN protein phylogenies (Figure 2 is arbitrarily midpoint rooted for clarity). As a result we cannot conclusively determine the origin or earliest representatives of the SodN enzyme. However, the abundance, diversity, and phylogenetic relationships of SodN sequences from Actinobacterota and Chloroflexota makes them prime candidates for the originators or earliest adopters of SodN. As previously highlighted, the Actinobacteriota clade encompasses organisms that live in a variety of environments, many of which are found in soil and sediment. The Chloroflexota phylum also contains incredible diversity in metabolism and environmental requirements. Notably, this phylum includes members that are thermophilic, heterotrophic, autotrophic, aerobic, and anaerobic (Islam et al., 2019). Representatives of the class Dehalococcoidia are well represented among the Chloroflexota members with *sodN*; these organisms are primarily found in terrestrial and marine sedimentary environments (Wasmund et al., 2014). Anaerolineae is the second most well represented Chloroflexota class in *sodN* diversity. This class contains members isolated from marine sediment, aquifers, and wastewater (Yamada & Sekiguchi, 2009). Although this class has been previously thought to include primarily obligate anaerobes that make a living from fermentation, several members contain genes for respiration (Hemp et al., 2015; Pace et al., 2015; Ward et al., 2015). The presence of NiSOD in diverse members of these two deeply rooted classes of Chloroflexota suggests the possibility that NiSOD was present in their last common ancestor. Molecular clock analyses have suggested that Anaerolineae and Dehalococcoidia diverged ~2.2 Gya (Shih et al., 2017), providing an indirect estimate for the antiquity of this enzyme family.

Comparison of organismal and SodN protein phylogenies can provide some limited insight into the timing of NiSOD evolution. If, as discussed above, the root of the NiSOD tree is near the branch separating the Chloroflexota and Actinobacterota clades (i.e. the midpoint root shown in Figure 2), this suggests that NiSOD was acquired by early members of these clades and has subsequently been vertically inherited into crown groups. Both Actinobacterota and Chloroflexota are phyla with obligately anaerobic basal lineages that are thought to have originated in Archean time before the Great Oxygenation Event, subsequently acquired aerobic respiration, and underwent radiation of class-level lineages during early Proterozoic time (Lewin et al., 2016; Shih et al., 2017). It therefore follows that NiSOD originated at about this time, most likely under low oxygen conditions in marine sediment or the water column. Though it may seem logical to constrain the timing of the evolution of NiSOD with the GOE and associated introduction of significant oxygen to marine environments, small amounts of oxygen can be produced by abiotic processes including photolysis or radiolysis of water—providing an evolutionary impetus for SOD evolution decoupled from biological oxygen production. The origin of NiSOD in late Archean or early Proterozoic time is further consistent with changing marine Ni availability through Earth history. Dissolved Ni concentrations have previously been hypothesized to be on the order of 400 nM in the Archaean ocean, decreasing to ~200nM by 2.5 Gya (Konhauser et al., 2009, 2015). These concentration estimates are more than 2 orders of magnitude greater than modern seawater concentrations (Sclater et al., 1976). Whether the ancestral NiSOD evolved in the marine water column or in underlying sediment, it almost certainly evolved in an environment where Ni exceeds modern marine concentrations.

There are several reasons why the presence *sodN* in a microbial genome may provide an advantage for organisms over its Fe-binding counterpart. One curious aspect of NiSOD is that it is the only metalloform whose metal cofactor does not react with superoxide as a hydrated ion. While the “ping-pong” mechanism of catalysis is shared by all SODs (i.e. higher valent metal center oxidize superoxide, lower valent metal center reduces superoxide in an alternating fashion), the redox potential of Ni(II/III) redox pair is tuned to the appropriate redox potential by its coordination within the enzyme (Herbst et al., 2009). Therefore, the presence of Ni and its availably is not influenced by O2 and ROS in the same way that Fe would be. The concentration of soluble Fe in seawater has decreased as a result of atmospheric oxygenation. Although estimates vary, Fe concentrations likely dropped from ~100 μM in Archaean seawater, to low μM or high nM concentrations in the Proterozoic, to the near 1 nM dissolved Fe seen in the open ocean today (Holland, 1984). Today, Fe can be a limiting nutrient for primary productivity in many surface ocean regions, stemming from its use as cofactor in many different essential enzymes (Behrenfeld et al., 1996; Behrenfeld & Kolber, 1999). The utilization of NiSOD over FeSOD under such iron limitation provides a competitive advance in its ability to reduce iron quotas.

NiSOD may offer an additional competitive advantage beyond reducing cellular Fe quotas. The superoxide-dismutating activity of both FeSOD and NiSOD are inhibited by excess hydrogen peroxide, one of the two primary products of the dismutation reaction (Sheng et al., 2014). However, the H_2_O_2_ inhibition of FeSOD and NiSOD work in fundamentally different ways. FeSOD inhibition by H_2_O_2_ occurs through irreversible peroxidative damage of the enzyme that proceeds through a Fenton-type reaction. Inhibition of NiSOD, conversely, operates by reducing Ni(III) to Ni(II) without damaging the enzyme itself (Herbst et al., 2009). This inhibition is reversible, and SOD catalytic activity is restored once hydrogen peroxide levels drop below the inhibition threshold. Similar to FeSOD, CuZnSOD may also be irreversibly inhibited by excess H_2_O_2_ (Gabbianelli et al., 2004). While MnSOD is not known to be inhibited by H_2_O_2_, its SOD activity is lessened slightly at higher concentrations of superoxide (Sheng et al., 2014), which, to our knowledge, has not been demonstrated for the other metalloforms.

In the surface ocean, superoxide and hydrogen peroxide production is wide-spread and results from a combination of light-dependent and light-independent reactions (Diaz et al., 2019; Sutherland et al., 2019; Vermilyea et al., 2010). Surface superoxide concentrations range from low pM to several nM in the open ocean, with higher concentrations typically found in productive surface waters (Sutherland, Grabb, et al., 2020; Sutherland, Wankel, et al., 2020). In some cases, superoxide may approach 100-200 nM in productive coral reef ecosystems (Diaz et al., 2016; Grabb et al., 2019). Hydrogen peroxide has a lifetime in seawater that is ~3 orders of magnitude longer than superoxide (i.e. typical H_2_O_2_ lifetime is hours to days), and correspondingly has concentrations that range from low nM in the deep ocean to as high as several hundred nM in productive sunlit surface water (Hopwood et al., 2017; Yuan & Shiller, 2005). The ability of NiSOD to catalyze superoxide dismutation at near diffusion limited rates and withstand elevated concentrations of hydrogen peroxide may make it uniquely suited for microbial fitness in the surface ocean with modern oxygen levels.

There is observational evidence to support the notion that not all SOD enzymes are perfect replacements for one another, even when cofactor concentrations are replete. For example, one study examined the growth of two strains of *Synechococcus* under low and high Ni concentrations to compare Ni requirements for NiSOD and urease (Dupont et al., 2008). In that study, one strain’s genome encoded for only NiSOD, while the other’s genome encoded for both NiSOD and CuZnSOD. The authors showed that under low Ni concentrations and adequate fixed nitrogen (resulting in reduced Ni quotas for urease), Cu/ZnSOD could not replace NiSOD activity. However, not all cyanobacteria have NiSOD; FeSOD is represented across cyanobacterial diversity. Even still, this observation demonstrates that there are enzyme characteristics beyond the metal cofactor identity that select for certain SOD metalloforms; these are not well known and should be the target of future study.

Interestingly, 9 of the 26 archaea that we identified with NiSOD genes also contain genes for proteorhodopsins. All nine of these organisms belong to the Poseidoniales clade, which is one of the most widespread archaeal groups in surface ocean waters (Rinke et al., 2019). The selective pressures experienced by these archaea, including nutrient availability, light levels, and ROS levels, mirrors those experienced by cyanobacteria such as *Synechococcus* and *Prochlorococcus*, suggesting that the acquisition and expression of *sodN* plays a key role in microbial success in the modern ocean.

## Summary and Conclusion

Using the Genome Taxonomy Database, we examined the phylogenetic relation of the nickel-binding metalloform of superoxide dismutase to better understand its evolutionary history and biogeography in the context of environmental constraints. We compared organismal phylogenies and NiSOD gene phylogenies to demonstrate that NiSOD has undergone multiple instances of inter-domain horizontal gene from bacteria to archaea. Actinobacteriota and Chloroflexota as two clades in which NiSOD is pervasive and in which the organismal phylogeny resembles that of the *sodN* gene. As such, these two clades are the most likely candidates for the evolutionary root of *sodN*. The distribution of NiSOD sequence diversity is consistent with one or more vertical migrations in biogeography, starting in soil or marine sediment and migrating to the photic zone. The acquisition of *sodN* as a means to mitigate iron-limitation is consistent with the biogeochemical history of the marine environment, however, other factors such as hardiness against damage by the product hydrogen peroxide may also contribute to its proliferation in the surface ocean.

## Acknowledgements

This work was funded by an Agouron Institute geobiology postdoctoral fellowship (KMS), the Simons Foundation (Grant Number 653687 to LMW), and Harvard University (CRC, DTJ).

